# Comparison of genomic diversity between single and pooled *Staphylococcus aureus* colonies isolated from human colonisation cultures

**DOI:** 10.1101/2023.06.14.544959

**Authors:** Vishnu Raghuram, Jessica J. Gunoskey, Katrina S. Hofstetter, Natasia F. Jacko, Margot J. Shumaker, Yi-Juan Hu, Timothy D. Read, Michael Z. David

**Affiliations:** Microbiology and Molecular Genetics Program, Graduate Division of Biological and Biomedical Sciences, Laney Graduate School, Emory University, Atlanta, Georgia, USA; Division of Infectious Diseases, Department of Medicine, Emory University, Atlanta, Georgia, USA; Division of Infectious Diseases, Department of Medicine, University of Pennsylvania, Philadelphia, PA, USA; Department of Biostatistics and Bioinformatics, Emory University, Atlanta, Georgia, USA

**Keywords:** Staphylococcus aureus, whole genome sequencing, asymptomatic carriage, adaptation, genetic diversity

## Abstract

The most common approach to sampling the bacterial populations within an infected or colonised host is to sequence genomes from a single colony obtained from a culture plate. However, it is recognized that this method does not capture the genetic diversity in the population. An alternative is to sequence a mixture containing multiple colonies (“pool-seq”), but this has the disadvantage that it is a non-homogeneous sample, making it difficult to perform specific experiments. We compared differences in measures of genetic diversity between eight single-colony isolates (singles) and pool-seq on a set of 2286 *S. aureus* culture samples. The samples were obtained by swabbing three body sites on 85 human participants quarterly for a year, who initially presented with a methicillin-resistant *S. aureus* skin and soft-tissue infection (SSTI). We compared parameters such as sequence quality, contamination, allele frequency, nucleotide diversity and pangenome diversity in each pool to the corresponding singles. Comparing singles from the same culture plate, we found that 18% of sample collections contained mixtures of multiple Multilocus sequence types (MLSTs or STs). We showed that pool-seq data alone could predict the presence of multi-ST populations with 95% accuracy. We also showed that pool-seq could be used to estimate the number of polymorphic sites in the population. Additionally, we found that the pool may contain clinically relevant genes such as antimicrobial resistance markers that may be missed when only examining singles. These results highlight the potential advantage of analysing genome sequences of total populations obtained from clinical cultures rather than single colonies.

**Graphical Abstract:** 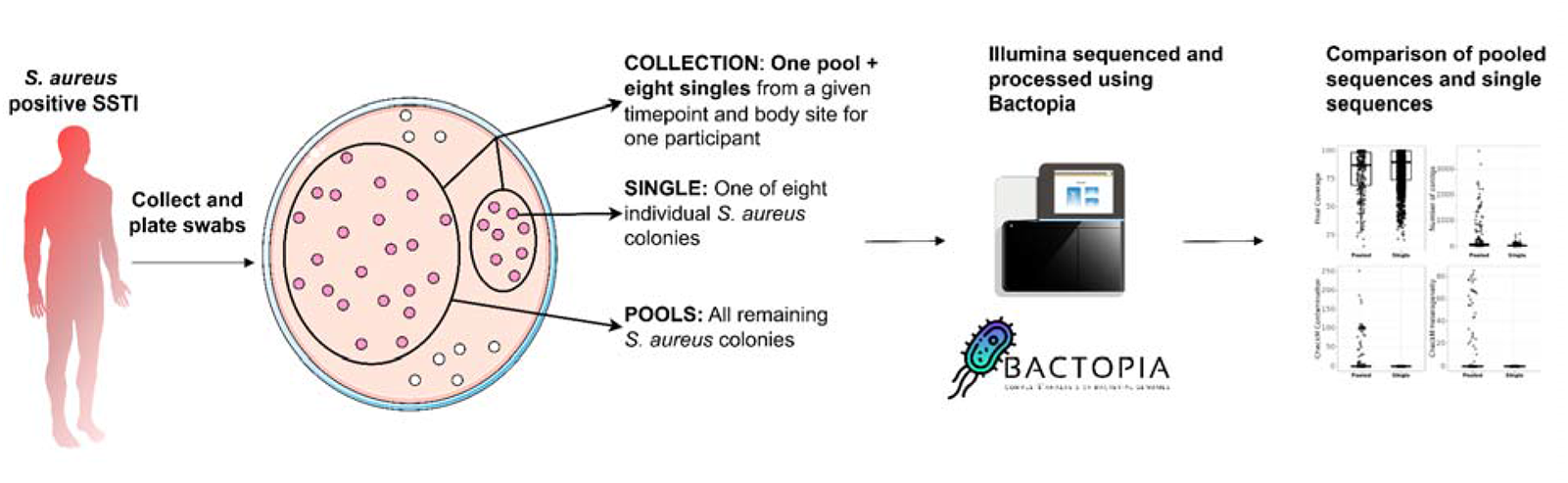

## Introduction

Large-scale Whole Genome Sequencing (WGS) of bacterial pathogen species offers hope for more accurate infectious disease surveillance and a better understanding of within-host evolution during human disease and asymptomatic colonisation (1–3). Typically, bacterial genomics-based surveillance studies sequence DNA isolated from single bacterial colonies cultured on selective media from each clinical sample tested (1,4–6). The single-colony bacterial culture can be further tested in the laboratory for phenotypes such as antibiotic resistance and toxicity.

However, individual colonies do not provide insight into the genetic diversity of the population of the species in the sample as there could be multiple strains present in the sample (7–9). Even if only one strain is present, there will be accumulated microdiversity between individual isolates that is roughly proportional to the duration between initial colonisation and time of sampling, assuming the absence of bottlenecks (5,10).

A few studies have undertaken sequencing multiple single colonies from clinical samples (9,11,12). This strategy can allow the comparison of phenotypes associated with intra-sample genetic variation and the construction of phylogenetic trees to trace the relationship between samples. However, costs rise linearly with each additional colony sequenced per sample, necessitating a cost-benefit analysis of how many colonies to sequence. Sequencing colonies from isolation plates of a single sample is an alternative approach (12–15). This is sometimes called “sweep sequencing”, “population sequencing” or “pool-seq”, and the latter term will be used here. In pool-seq, multiple colonies from the same species can be sampled genetically at the same economic cost as sequencing an individual isolate. Pool-seq has generally been found to be reliable in measuring sequence variation and allele frequency (11,16). However, the disadvantages of this method are the perceived complexity of the bioinformatic analysis and the complications of assessing phenotypic characteristics of the population of bacterial clones, such as antibiotic resistance, when these assays typically require clonally purified single colonies.

Single-isolate sequencing is a convenient sampling strategy based on the assumption that strain mixtures are rare and capturing within-strain microdiversity is not worth the additional expense of sequencing. While the cost of raw sequence production has steadily declined, costs of labour and infrastructure, such as DNA extraction, library preparation, physical sample storage, and bioinformatic analysis, have not scaled down at the same rate (17). There has also been little analysis of what the increased sequencing and storage costs from sampling multiple colonies or pools yield over single colonies. However, pool-seq can still provide insights into the natural history of even well-studied pathogens, and inform us about the fate of adaptations that enhance virulence and antibiotic-resistance (1,18–21). Therefore, optimising sampling strategies and genomic workflow design is essential to minimise the number of samples processed while maximising the information obtained from each clinical sample.

In this work, we use samples from an ongoing study of *Staphylococcus aureus* colonisation on humans to compare the three strategies outlined above: single-isolate sequencing, sequencing collections of multiple single colonies, and pool-seq. *S. aureus* is a ubiquitous nosocomial pathogen prevalent worldwide, causing invasive disease syndromes such as bacteremia, endocarditis, and osteomyelitis (22,23). Like other prominent pathogens, WGS has significantly improved *S. aureus* epidemiologic studies, and our ability to track the spread of antibiotic resistance and virulence across populations (2,24–28). Here, we used samples from human participants who had an index methicillin-resistant *S. aureus* (MRSA) skin and soft tissue infection (SSTI) as part of an ongoing study, SEMAPHORE (Study of the Evolution of MRSA, Antibiotics and Persistence Having the Outcome of Recurrence). The study was designed to examine the clinical and demographic characteristics of the participants, and the genomes of the colonising *S. aureus* to identify factors associated with recurrent skin infections. However, for this paper, we focused on the relationship between the pool-seq and collections of single isolate genome sequencing. We first quantified the amount of variation within the collections of single genome and pool-seq and then investigated three specific questions: 1) Can the pool-seq data identify clonal *S. aureus* populations (comprising a single ST) from mixtures of diverse lineages?; 2) Can pool-seq data be used to estimate the number of sites within mono-ST populations undergoing polymorphisms?; 3) Was pool-seq more sensitive in detecting antimicrobial resistance (AMR) genes than sequencing single clones?

## Results

Samples from the SEMAPHORE study were plated on CHROMAgar *Staphylococcus aureu*s and a “collection” of eight individual *S. aureus* colonies (“singles”) was obtained (**Fig 1B**). The remaining *S. aureus* colonies on the plate were pooled and sequenced, hereafter referred to as “pools” or “pool-seq”. The collective sequencing data obtained from all eight singles for each pool were referred to as “expected pools”. Similarly, sequencing data sampled from two random singles and four random singles, respectively, were combined to generate “downsampled pools”. The SEMAPHORE data used in this work had 85 participants with 254 samples (254 pools and 254 collections of eight singles – 2032 singles total) (**Fig 1A**). All FASTQ files (pool-seq and singles) were capped to 100x *S. aureus* genome coverage.

**Fig 1:**
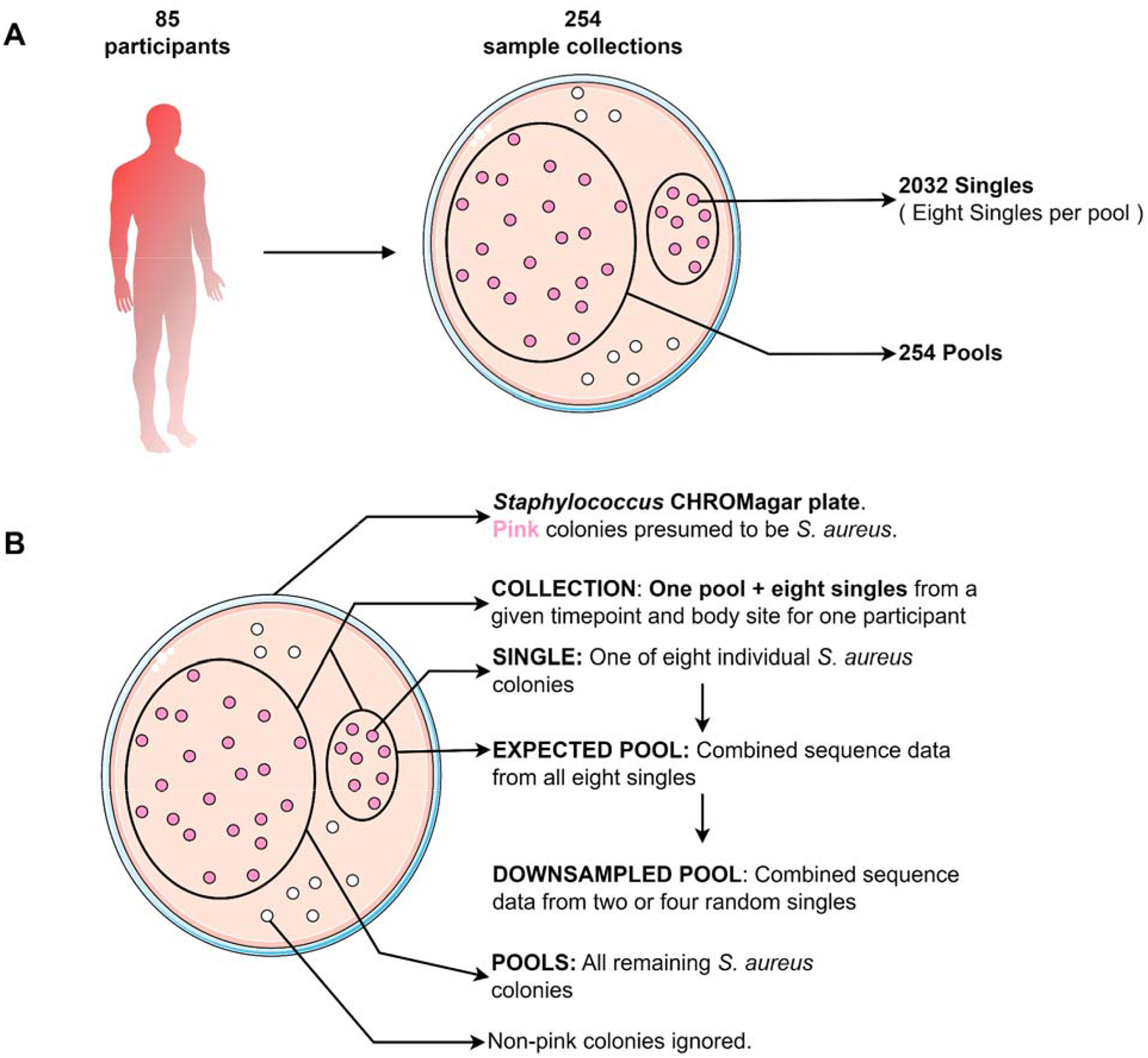
Schematic representation of colony collection strategy, names, and descriptions of isolate groups analysed in this study. (**A**): Diagram depicting the number of samples in this study. From 85 study participants, we collected a total of 254 culture samples. For each collection, we obtained 8 single colonies, and pooled the remaining colonies on the plate. (B): Diagram describing the terminology for specific isolate groups used in this study.

### 82% of collections have only one Multilocus Sequence Type (MLST)

Upon analysing the singles alone, we found a wide range of SNP distances between singles from the same collection, with a minimum of 0 and a maximum of 15,315. For 241 out of 254 collections (∼95%), the maximum SNP distance between any two pairs of isolates was < 100 (**Fig 2A**), suggesting that most collections of eight singles comprised only closely related isolates. However, 12 collections of singles (5%) showed clear signs of mixed strains, with a maximum SNP distance > 4000. When we compared the MLST amongst the singles for each collection, all 12 of these collections had at least one isolate that was a different ST from the remaining. This showed that comparing STs and pairwise SNP distances between multiple singles within collections could identify potential mixed infections, as mono-ST collections had lower maximum pairwise SNP distances. Although 59 STs were identified in total, 51% of singles (1051/2032) belonged to ST8 and ST5 (**Fig 2B, D**).

When we included both pools and singles in the ST analysis, we found that for 209/254 collections (∼82%) the ST types for the eight singles and the pool were identical, suggesting they were mono-ST collections. In the remaining 45 collections (∼18%) either at least one single or the pool had a different ST (**Fig 2B, C**). This includes the 12 aforementioned collections where we detected multi-ST collections based on singles alone. This means we found more potential multi-ST collections when we included the pools in our analysis. For 15 out of these 45 collections, the ST of the pools were untypeable due to the presence of multiple alleles for the same gene, confirming that these were multi-ST collections based on observing the pool alone. This means that for 30 collections, we needed to refer to the ST types from both the pool-seq and from the corresponding eight singles to infer whether the sample was multi-ST.

We observed no significant differences in the occurrence of multi-ST pools across the different timepoints, body sites and culturing methods (Chi-squared test, p>0.01). These data suggested that a given collection usually had a low level of *S. aureus* diversity and that we can find ST mixtures by comparing SNP distances and ST types within collections of single colonies.

**Fig 2:**
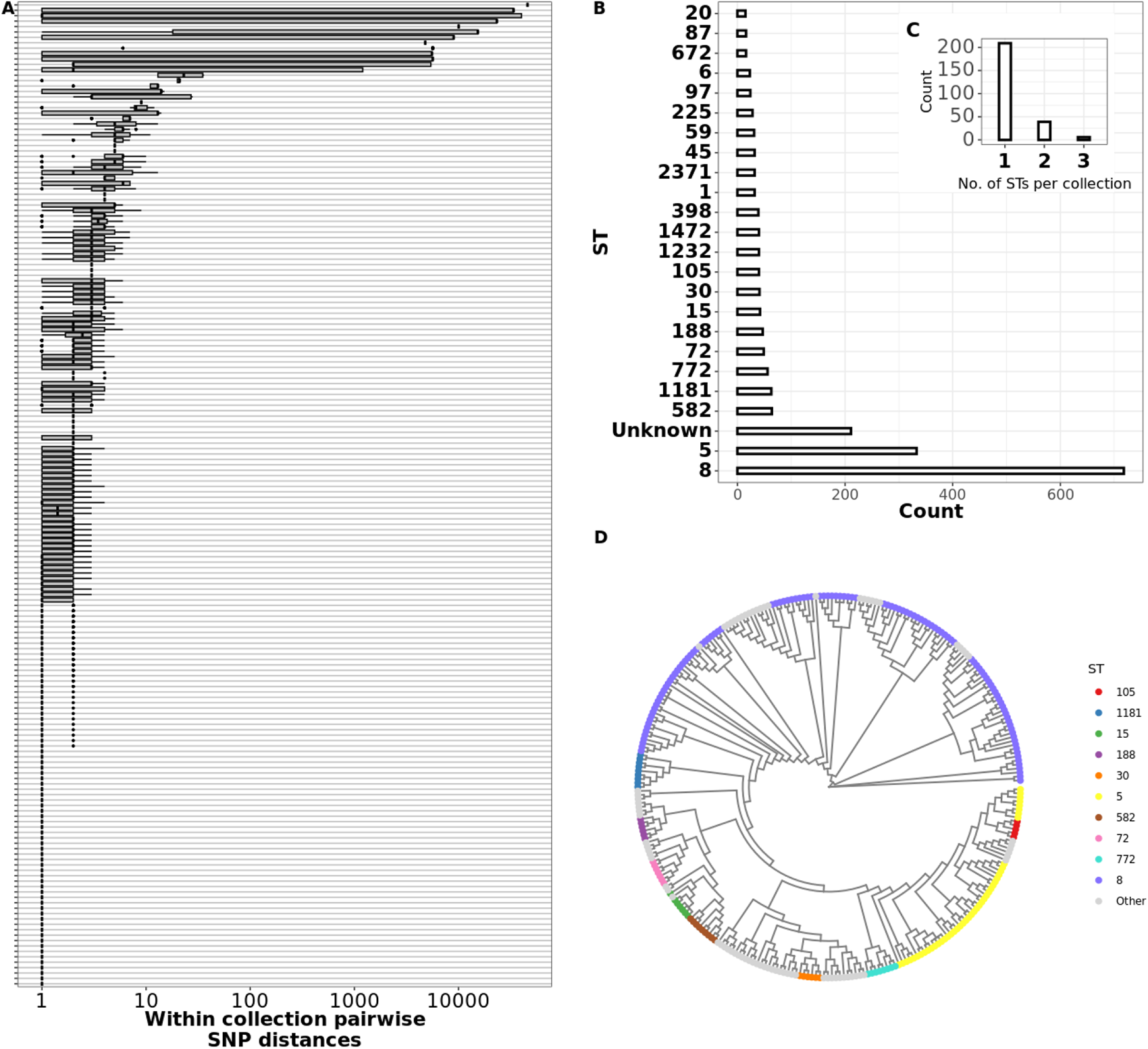
Pairwise SNP distance between and within collections. (**A**) Boxplots showing per-collection SNP distance distributions. For each collection shown in the y-axis, the x-axis shows the corresponding distribution of core genome SNP distances in log scale. Black vertical lines show the median SNP distances and boxes show the interquartile range. Whiskers represent values up to 1.5 times the first or third quartile. Black dots represent outliers beyond the whiskers range. (**B**) Barplot showing number of genomes per ST. Multilocus Sequence Typing was performed by the software tool mlst (see methods). x-axis shows the number of isolates assigned to the corresponding ST shown in the y-axis. (**C**) Bar plot showing number of STs detected per participant. Multilocus Sequence Typing was performed for all eight singles from a participant and the number of unique STs detected per participant was plotted. (**D**) Maximum likelihood phylogeny representing at least one isolate from all collections. All non-identical genomes from each collection were aligned by snippy and a core genome phylogeny was constructed using fasttree (see Methods). Tree tips are coloured by ST, only top 10 most frequent STs are shown, and remaining are grouped into “Other”.

### Pool-seq samples with elevated average minor allele frequency, elevated number of contigs, higher nucleotide diversity and untypable MLST were associated with strain mixtures

As mentioned previously, for 30 collections, we were not able to confirm the presence of a multi-ST population based on the ST assignment in the pool-seq alone. Therefore, we examined what features of the pool-seq data could be used to assess whether there was one ST present in the pool, or a mixture of STs by comparing the 254 pool-seq samples to their cognate collections of singles. We focused on four measures: assembly quality, nucleotide diversity, gene number and minor allele frequency (MAF).

Sequencing reads from both singles and pool-seq were processed identically using the Bactopia pipeline with the same quality control parameters (29). Both single and pool-seq reads had a final average quality score of 36.3 (Welch’s t-test p > 0.01). We expected the genome assemblies (generated using the SKESA assembler (30)) from single colonies to be higher quality than pool-seq, as the latter may contain multiple *S. aureus* strains and possibly contaminating species from the culture plate. We evaluated assembly quality using CheckM and QUAST (31,32), observing that, while most pools and singles had comparable coverage (**Fig 3A**, Wilcoxon p>0.01, effect size=0.052), pools had higher number of contigs (**Fig 3B**, Wilcoxon p < 0.01, effect size = 0.20), higher heterogeneity (**Fig 3C**, Wilcoxon p < 0.01, effect size 0.347), and contamination scores (**Fig 3C**, Wilcoxon p < 0.01, effect size = 0.239). 32 out of 224 pools (14%) had more than 200 contigs in contrast to only 5 out of 1792 singles (0.2%). The CheckM heterogeneity score indicated the source of the contamination – a heterogeneity score < 50% indicates that the source of contamination is phylogenetically distant and vice versa (30). While all singles had contamination and heterogeneity scores of 0, the pools ranged from low heterogeneity contamination to high heterogeneity contamination (**Fig 3C**). 7 pools (3%) had a heterogeneity score > 50 with a contamination score >10, suggesting they were contaminated by phylogenetically similar sources. 15 pools (6%) had a heterogeneity score < 50 with a contamination score >10, suggesting they are contaminated by phylogenetically distant sources. Overall, these results suggested that genome assembly quality can be useful for assessing population heterogeneity of pool-seq.

**Fig 3:**
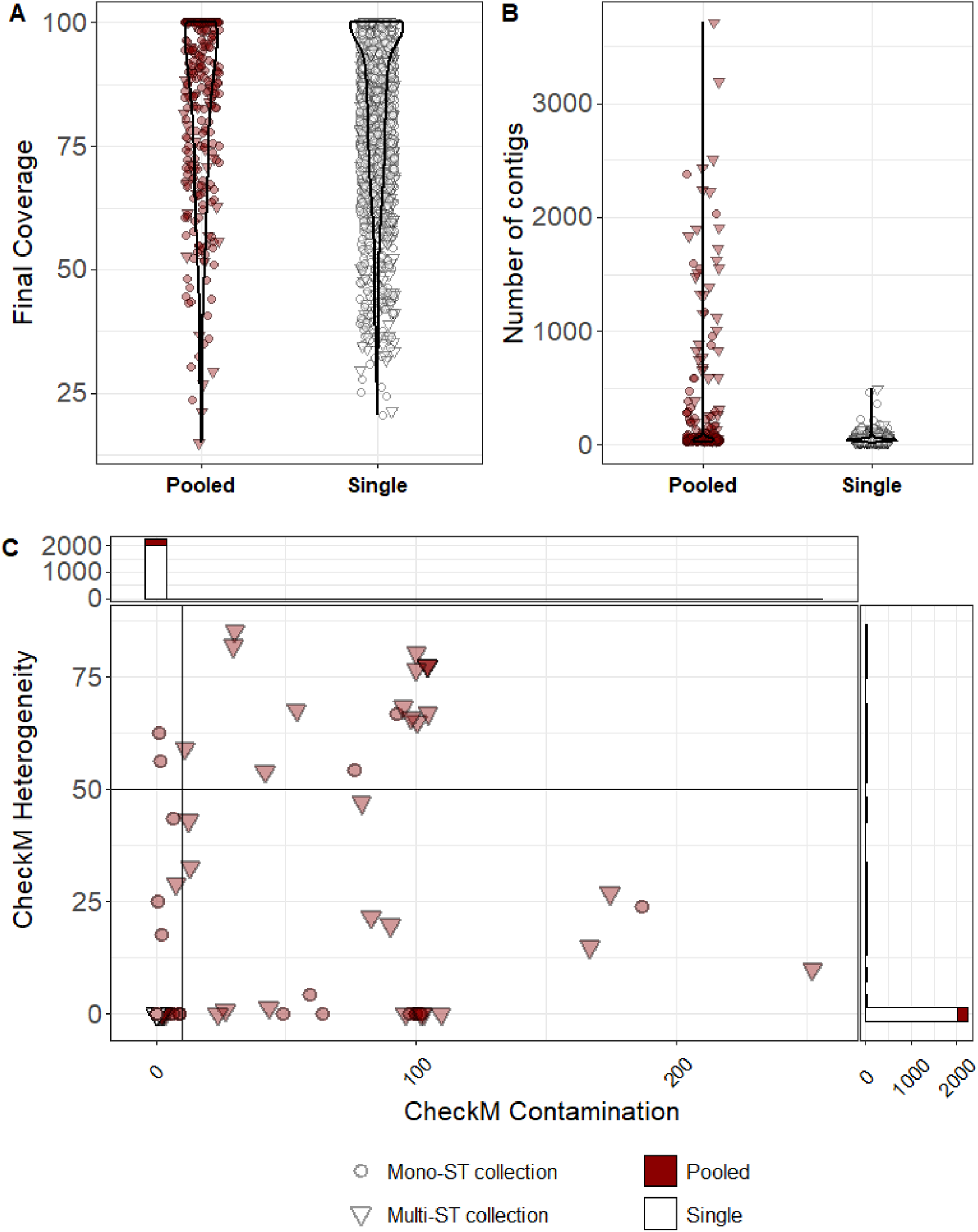
Assembly quality can be used to assess population heterogeneity. **(A) There was no significant difference in the assembly coverage between pools and singles**. Violin plot showing distribution of assembly coverage between pools and singles. Assembly coverage for each pool and single was calculated by Bactopia against an auto-chosen reference (see methods). Circles indicate mono-ST collections and triangles indicate multi-ST collections. **(B) Pool assemblies were more likely to have a higher number of contigs than single assemblies.** Violin plot showing distribution of number of assembly contigs in pools and singles. Pooled samples were processed identically to singles with Bactopia using SPAdes. Circles indicate mono-ST collections and triangles indicate multi-ST collections. **(C) Pooled samples have varying sources of contamination while singles are pure**. CheckM contamination and heterogeneity scores showed that all single colonies have no contamination while 6% of pools are contaminated by phylogenetically distant sources and 3% of pools are contaminated by phylogenetically similar sources. The blue line marks a heterogeneity score of 50 below which the source of contamination is considered phylogenetically distant and vice versa. Circles indicate mono-ST collections and triangles indicate multi-ST collections.

Allele frequency (AF), i.e., the fraction of reads piled up over a particular variant position is a useful metric of genomic heterogeneity. We mapped the pooled sequences against the closest complete *S. aureus* reference genome (see methods) to obtain the variant sites. The eight singles were also aligned to the same reference as the corresponding pool. For each sample in a collection, we calculated the average minor allele frequency, i.e., the total number of variant sites divided by the sum of all minor allele frequencies (MAF). If all reads mapped to only the reference or only the alternate alleles, the average MAF would always be zero, as would be expected from ideal single pure cultures. We plotted the average MAF against the total number of variant sites for 254 pools (**Fig 4A**). We split the plot into four quadrants based on two parameters – the number of variants cutoff of 2800 sites (or 0.1% of the *S. aureus* genome) suggesting only few variant sites, and average MAF cutoff of 0.05 below which we deemed the sample as having no minor alleles. 223 pools (∼88%) had a total number of variant sites less than 0.1 % of the *S. aureus* genome (**Fig 4A left quadrant**). 55 out of these 223 had an average MAF < 0.05 (**Fig 4A bottom left quadrant**), suggesting highly homogeneous samples. In contrast, there were 14 pool-seqs with more than 2800 variants (∼0.1%) of MAF > 0.05 (top right quadrant in **Fig 4A**). Pool-seqs in the bottom right quadrant (average MAF < 0.05 with > 2800 variants) have a large number of variants because they are distant from the reference sequence used for variant calling; however, their low average MAF (< 0.05) indicates they are still pure samples (see methods for how reference sequences were chosen). To assign a diversity score based on MAFs, we used the product of the total number of variants and the average MAF, which we termed the “MAF Index”. The MAF Index was higher for samples with both a large number of variants compared to reference as well as a high average MAF (i.e., top right quadrant in **Fig 4A**). The MAF index of mono-ST pools was not significantly greater than the MAF index of singles (Welch’s t test, p>0.01). However, within the pools, the multi-ST pools had significantly greater MAF index than the mono-ST pools (Welch’s t test, p< 0.01).

AFs are measured based on whether or not a given position in a read maps to the reference, but our calculations did not take into account the possibility of multiple alternate alleles (which we assumed to be very rare given that the number of variants was only a small percentage of the chromosome). Therefore, we also calculated intra-sample nucleotide diversity between the true pooled sequences and our expected pools as an analogous method for measuring genomic heterogeneity. Using the InStrain software (33), we estimated nucleotide diversity (π), which is π for each position, 1 minus the sum of the frequency of each base squared (1 – [(frequency of A)∧2 + (frequency of C)∧2 + (frequency of G)∧2 + (frequency of T)∧2]). This value was then averaged across the whole genome. We used InStrain to measure the average across our π singles, downsampled pools (four and two colony pools), expected pools and true pools. The π value was significantly greater in pools compared to singles (Welch’s test, p < 0.01). However, in 216 pools (96%), the diversity observed in the true pools was less than the expected diversity observed from a simulated mixture of two *S. aureus* isolates 30,000 SNPs apart in a 99:1 ratio (**Fig 4B LEFT** black horizontal line; see Methods). This analysis provided evidence that most pools comprised only single strains. Moreover, similar to average MAF, π of multi-ST pools were significantly greater than π of mono-ST pools (Welch’s test, p < 0.01). In cases where there was an increased diversity value for our expected pools or downsampled pools compared to our true pools, we may have overestimated the diversity of our true pool by assuming the single colonies were present in equal abundance. Alternatively, since we pooled only the remaining colonies on the plate after picking singles, we may have reduced some diversity from the pool by removing certain single colonies. In the 12 cases where the true pools had a diversity value greater than the corresponding expected pools (5%), the eight colonies sampled did not capture the entire diversity of the pool.

**Fig 4:**
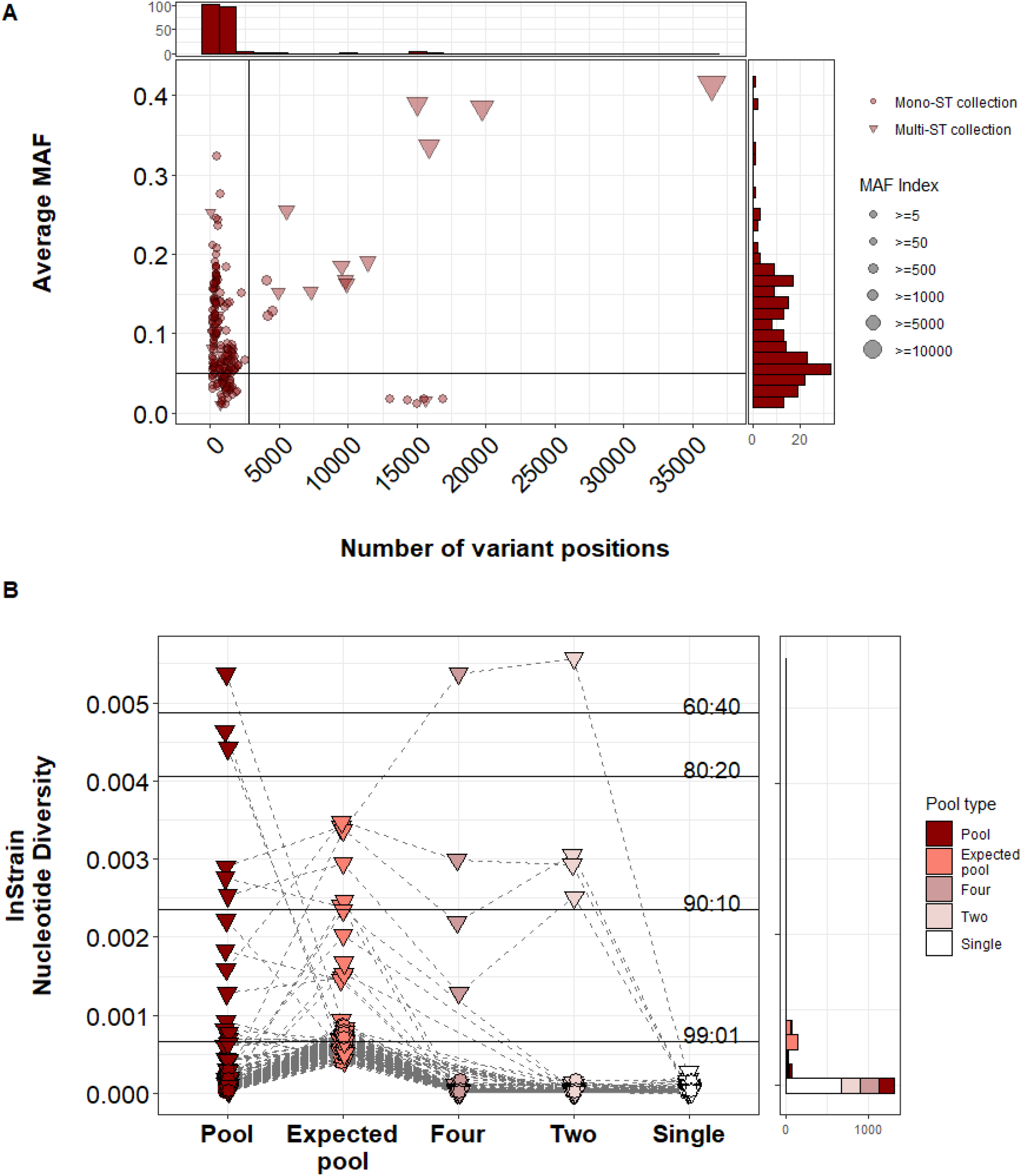
**(A) The MAF index could be used to assess multi-ST pools.** Dot plot depicting the number of variant positions and the average MAF for mono-ST (circles) and multi-ST (triangle) pools. The x-axis indicates the number of variant positions compared to a reference. The y-axis indicates the average minor allele frequency (MAF). The average MAF was calculated by summing the MAFs of all intermediate alleles and dividing by the total number of variant positions. Red dots correspond to mono-ST pools and triangles correspond to multi-st pools. The black horizontal line indicates an average MAF of 0.1. The black vertical line indicates 0.1% of the *S. aureus* genome (2800 sites). The frequency of the dots at their corresponding x and y positions are indicated by the histogram above the x- and to the right of the y-axis. **(B) Average nucleotide diversity suggested most pools comprise single strains. 94% of pools had nucleotide diversity less than a theoretical 99:1 mixture of two strains.** LEFT: Dots and colours indicate average nucleotide diversity value for each pool, expected pool (reads from eight singles combined in equal proportions), downsampled pools (i.e., reads from four and two random singles combined in equal proportions) and all 2032 singles. Grey dashed lines connect corresponding samples. Black solid horizontal lines indicate the average nucleotide diversity value for in-silico mixtures of two *S. aureus* genomes 30,000 SNPs apart. The ratio of each mixture is indicated over each solid black line. The frequency of the dots at their corresponding x positions are indicated by the histogram to the right.

We have shown that the number of contigs, contamination, minor allele frequencies, and nucleotide diversity are significantly different between pools and singles (**Fig3**, **Fig4**). Next, we wanted to measure the magnitude of these parameters’ contribution to the variability between pools and singles.

We performed principal component analysis (PCA) for five different parameters we measured (CheckM contamination, CheckM heterogeneity, MAF Index, Nucleotide diversity, and the number of contigs). PC1 and PC2 explained ∼78% of the total variance (**Fig S1A**). All five parameters had positive loadings in PC1 (>0.4) and the CheckM contamination score had positive loadings in PC2 (>0.6) (**Table S1**). This result suggested that the deviation of some pools from the singles was mainly due to contamination (PC1) and allelic variation (PC2).

We also performed an all vs. all Pearson’s correlation across the five different parameters mentioned above (**Fig S1B**). We found that CheckM contamination, number of contigs, and CheckM heterogeneity were positively correlated with each other, suggesting that contamination reduced the assembly quality (larger number of contigs). However, these three parameters did not have a high positive correlation with the MAF index or with nucleotide diversity. This showed that contamination and allelic diversity can independently drive heterogeneity in the pool, and that pooling multiple colonies may impact sequencing and assembly quality regardless of intra-species diversity.

From our analysis thus far, we have shown that there are pools that provide information similar to pure singles, and there are true strain mixtures. As we mentioned earlier (**Fig 2**), detecting multiple MLST types in the pool or measuring pairwise SNP distances between singles from within a collection are a reliable way to ascertain true mixtures. However, when the MLST calls are unreliable (unassigned types/undetectable alleles) or hypothetically if we did not have single colonies, alternative methods would be required. Therefore, we wanted to test whether the above mentioned parameters (Number of contigs, CheckM contamination, CheckM heterogeneity, MAF index, nucleotide diversity) could serve as predictors for mixed pools and homogeneous pools.

We performed a logistic regression using the number of contigs, CheckM contamination and heterogeneity, the MAF index, and nucleotide diversity as predictor variables to calculate the probability that a given pool is mixed (see Methods). Here, we defined a mixed pool or multi-ST pool as a pool with multiple ST calls, or a pool corresponding to a collection of singles where the maximum pairwise SNP distance is > 2800 (0.1% of the *S. aureus* genome). Our logistic regression model showed strong predictive ability with a McFadden R^2^ of 0.59, sensitivity of 1, specificity of 0.94, and a receiver operating characteristic (ROC) curve with an area 0.86 (**Fig S1C**). The maximum variance inflation factor (VIF) across our predictor variables was < 2.3 indicating low multicollinearity. Overall, these results showed that by using information from multiple statistics, the pool-seq data alone was sufficient to predict the presence of multi-strain populations with high accuracy.

**Fig S1:**
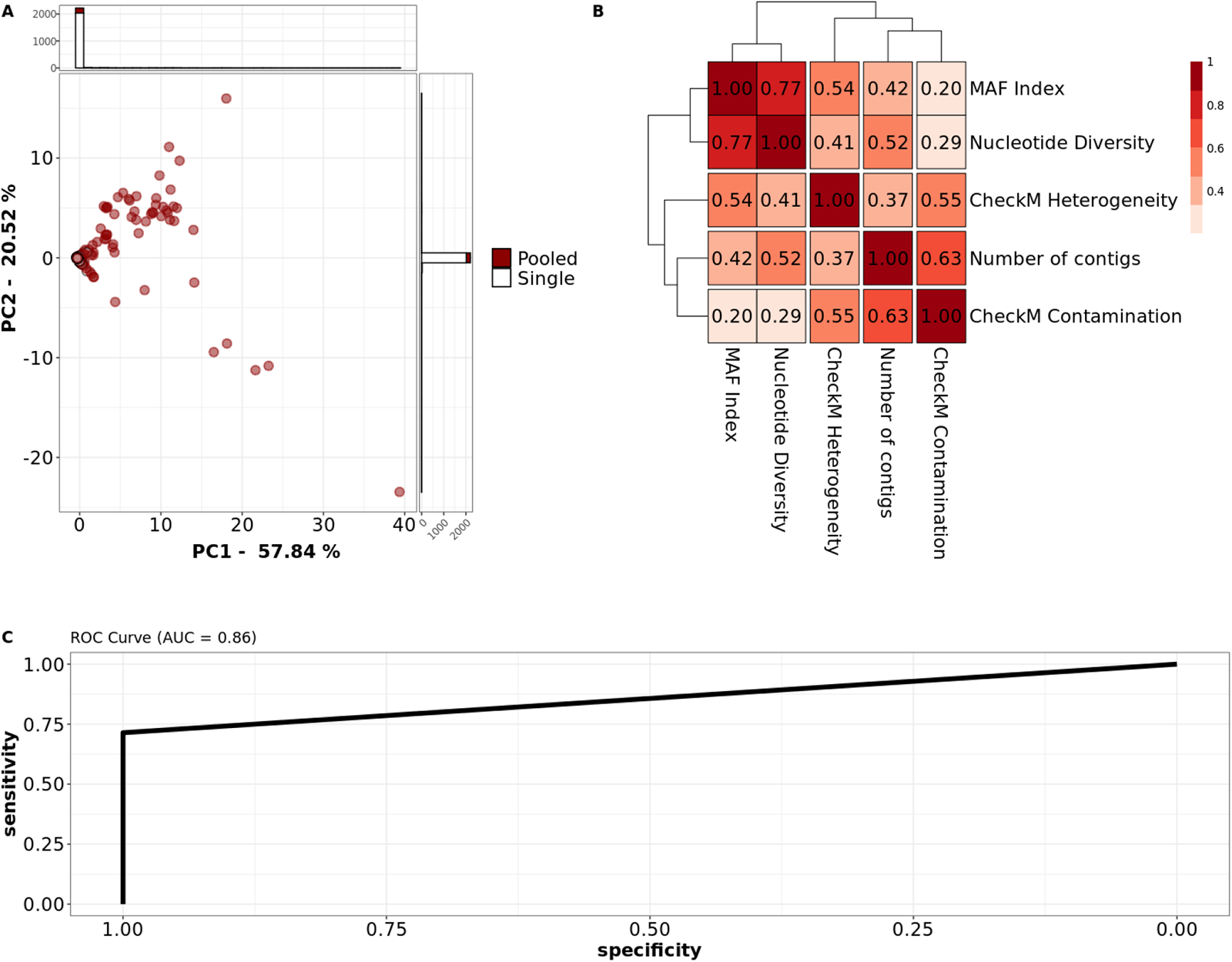
Variation in pools was primarily driven by contamination and allelic diversity. (**A**) PCA loading plot for principal components (PC) 1 (x-axis) and 2 (y-axis) explaining 70% of the total variance. White dots represent singles and red dots represent pools. 256 pools and 2032 singles were used for PCA. The density of the dots at their corresponding x and y positions are indicated by the histogram above and to the right of the plot respectively. The variance explained by each PC is indicated in the corresponding axis labels. (**B**) Pearson’s correlation coefficient matrix across five different diversity metrics. Each square indicates the Pearson *r* for comparing the corresponding parameters as labelled in the x and y axis. Scale indicates Pearson’s *r* (Darker = higher *r*) (**C**) Receiver operating characteristic (ROC) curve of the logistic model predicting multi-ST pools from parameters in **A** and **B**. Area under the curve (AUC) = 0.86.

**Table S1.** Summary of all five principal components (PC1 -– PC5) for five parameters used in Fig S1. All 254 pools and 2032 singles were used for principal component analysis.

|  | **PC1** | **PC2** | **PC3** | **PC4** | **PC5** |
| --- | --- | --- | --- | --- | --- |
| **MAF Index** | 0.462809 | -0.5268234 | 0.1029145 | -0.186782 | -0.6802838 |
| **Number of contigs** | 0.4541359 | 0.288277 | -0.5902644 | -0.5812948 | 0.1560169 |
| **CheckM Contamination** | 0.4020166 | 0.6543029 | 0.0179086 | 0.5201767 | -0.3733174 |
| **CheckM Heterogeneity** | 0.4418238 | 0.1308287 | 0.7578247 | -0.2563222 | 0.3842866 |
| **Nucleotide Diversity** | 0.4719563 | -0.4405963 | -0.2576384 | 0.5393737 | 0.4752164 |

### Numbers of variants in pool-seq and eight singles from the same sample are correlated but pool-seq had greater number

One of the advantages of pool-seq over groups of singles is the potential to discover mutant subpopulations that may be missing in samples of individual clones. We measured the number of variant positions that were shared between the pool and at least one of the eight singles. For each collection, we calculated the number of variant positions seen both in the pool and in at least one of the corresponding eight singles as a fraction of the total number of variant positions observed. For analysing variants in the singles and the expected and downsampled pools that were built from singles, we only considered sites with an AF > 0.95. We found that 152 collections out of 254 (∼60%) had shared variant fraction >0.5, meaning more than half the variants found in each pool and the corresponding singles were identical for 60% of our samples **(Fig S2A)**. Curiously, we observed 30 collections (∼12%) having a shared fraction < 0.05. This is what would be expected if these singles and pool-seq were not from the same sample **(Fig S2B)**. These collections may have been mis-sampled, and we opted not to use them for further comparisons of pools and singles within the same sample. This brought down our total number of collections from 254 to 224. Out of the 224 pool-seq samples, 204 were cases where the pool-seq and all eight singles had the same sequence type.

We found that the number of variants found in the pools was greater than the combined number of variants from the eight singles in 178 out of 224 samples (∼79%). This was as expected as the pools should more often contain more individual isolates than the collections.

To illustrate this point further we compared the number of variants detected in the pools against eight singles combined (expected pool), four random singles and two random singles combined (downsampled pools) and one random single. This was done to answer the question – How many variants would we have seen if we had sampled only eight colonies/only four/only two/only one? We considered a variant present in the expected or downsampled pools if it was present in at least one of the sampled singles at an AF > 0.95.

We found that 198 pools (∼88%) captured more than 75% of all the variants in a collection (**Fig 5A**). This was significantly greater than the number of expected pools (129 pools or 56%) that captured a fraction of variants > 0.75 (Kolmogorov Smirnov *p* < 0.01). If we had sampled only one single colony for each collection, only 39% of the singles would have captured a fraction of variants > 0.75 (**Fig 5A**)-“one colony”).

Though the number of variants observed in the pools were usually greater than in the singles, we found that the more singles a variant was present in, the more likely we were to detect the same variant in the pool. We counted the number of singles each variant was present in, plotted it against the AF of the same variant in the pool, and found a strong positive correlation (Pearson *r* = 0.83) (**Fig 5B**).

**Figure S2:**
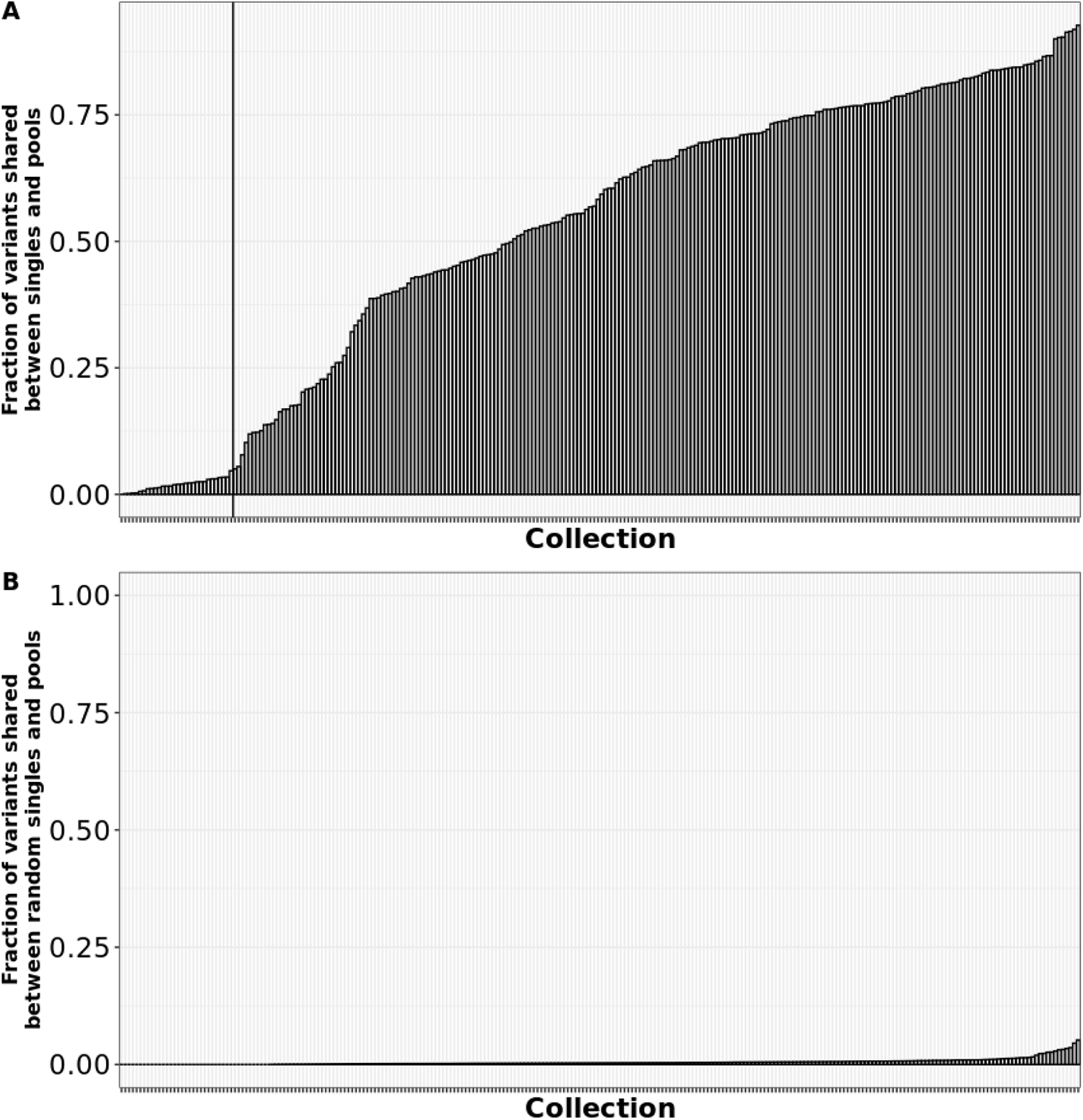
Collections with <5% of their total variants shared between pools and singles were discarded. **(A) Number of shared allelic sites revealed differences in the amount of diversity captured by single colonies and pools.** **Each bar indicates a collection, and the height of the bar indicates the fraction of variants shared between the pools and at least one of the eight corresponding singles.** Black vertical line indicates the threshold for shared fraction below which the singles and pools are not from the same sample (< 5% of variants shared) **(B) Expected fraction of allelic sites shared between a pool and a random collection of eight singles**. Each bar indicates a collection and the height of the bar indicates the fraction of variants shared between the pool and at least one of eight singles from a random other collection. The maximum observed fraction did not exceed ∼5% after 10 repetitions.

**Fig 5:**
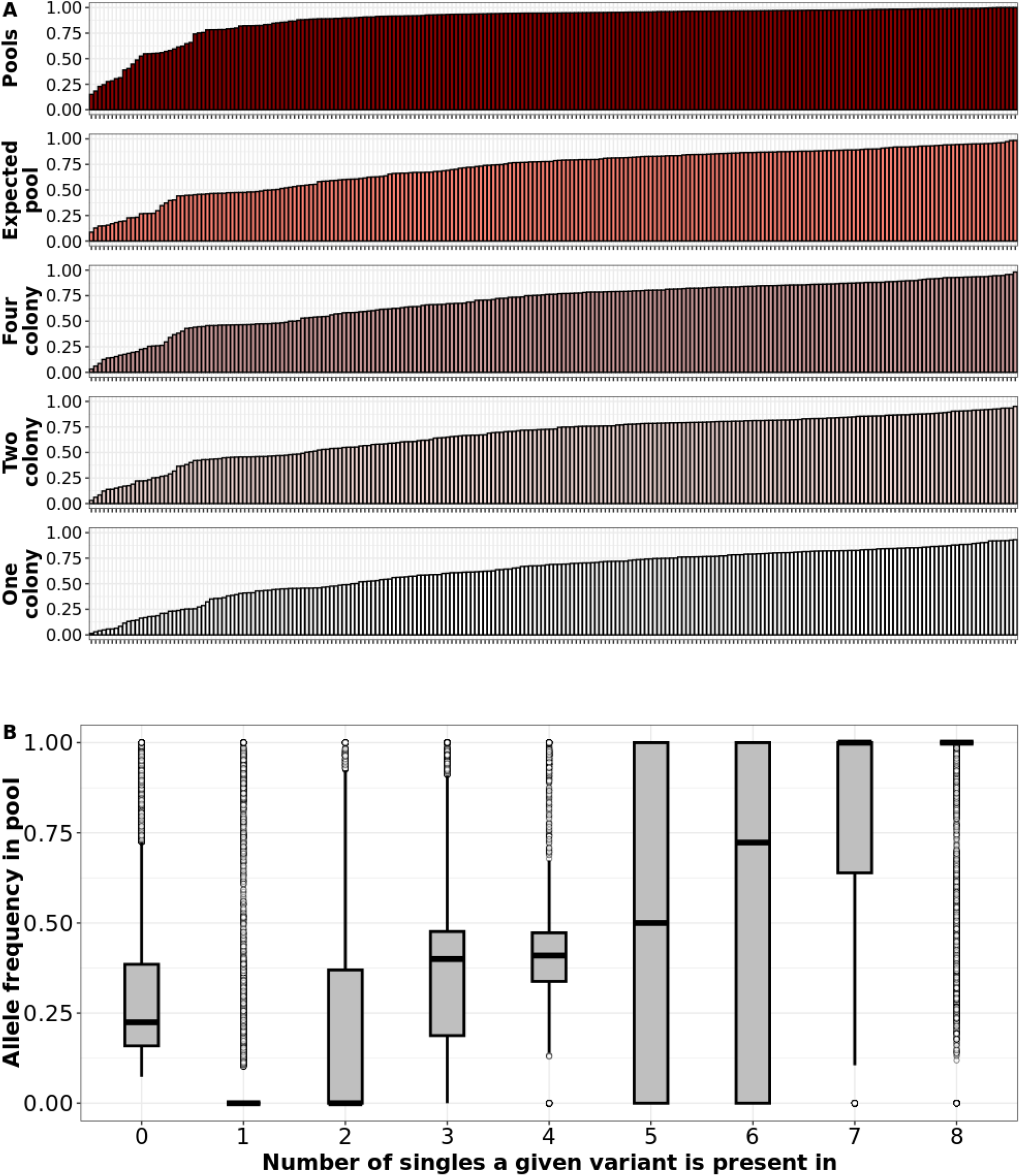
**(A) Pools captured more variants than eight single colonies combined.** Each bar indicates a collection, and the height of the bar indicates the fraction of variants found in the corresponding sample group (Pools, expected pools, four colony pools, two colony pools, single colony) to the total number of variants found in all samples in the collection (Pool plus all eight singles). For example, a bar with height 0.25 in the fifth row (**One colony**) shows that if one random single colony was examined from the specific collection corresponding to the bar, we would find 50% of the total number of variants found in the collection (Pool plus all eight singles). Bars for each sample group are ordered by lowest to highest. A value of one indicates 100% of the variants found in both the pools and all eight singles combined are represented in the sample group. **(B) Allele frequencies in the pool were proportional to the number of singles the variant was detected in.** Boxplots showing allele frequencies of variants detected in zero singles up to eight singles. Allele frequency of each variant found in the pool increased as the variant was found in more colonies in the corresponding singles. Boxes show the interquartile range and whiskers represent values up to 1.5 times the first or third quartile. White dots represent outliers beyond the whiskers range. Black horizontal line in each boxplot indicates the mean.

### Numbers of segregating sites in pools and singles from the same sample are positively correlated

As a bacterial population expands from an introduction event, mutations accumulate as a function of time (34,35). Subpopulations can segregate from the parent population by accumulating nucleotide variants at different sites across the genome and the total diversity across different subpopulations can be altered by bottlenecks and selective sweeps (8). The number of segregating sites (or within-population polymorphic sites) can therefore be an important indicator of the demographic history of the population, and it would be useful to know how well the pool-seq data could be used to estimate this value. Because we compared pools and singles to a common reference, a certain number of variants were likely fixed in the ancestor of the population. We expected these to have an AF of 1 or close to 1 (0.95 or greater) in both the pools and singles and filtered them out. We also filtered out samples where the ST of the pools and collection did not match the ST of the auto-chosen reference, as this would lead to an elevated number of variants. We found a moderate positive correlation between the number of segregation sites in the true pools and expected pools of the 198 samples that had matched ST across the pools, singles, and the reference (**Fig 6A**; Pearson *r* = 0.352). The number of segregating sites in the collection ranged from 8 to 1658. While the number of segregating sites was comparable between the true and expected pools, we also wanted to measure whether the proportion of the variants in the singles could reliably predict the proportion of the same variants in the pools. For each collection of sequences, we plotted the AF of the segregating sites observed in both the expected pool sequences and the true pooled sequences and calculated Pearson’s coefficient (*r*). If the AF of a variant present in an expected pool was equal to the AF of the same variant present in the true pool, we inferred that the proportion of the variant in the two populations was comparable. In other words, the variant frequencies in the eight singles combined (for example, if variant present in seven out of eight singles, AF = 0.875, if variant present in six out of eight singles, AF = 0.7, and so on) was equivalent to the variant frequencies in the pool. In contrast, if the AF of variants between the expected and true pools were not comparable, the pool-seq was significantly different from the expected pools. The distribution of *r* values indicated only 82 collections (41%) with *r* > 0.5 (**Fig 6B**). This result showed that in only less than half of our collections with matched ST, the proportion of the variants in the singles are positive predictors of the proportion of the same variant in the pool.

**Fig 6:**
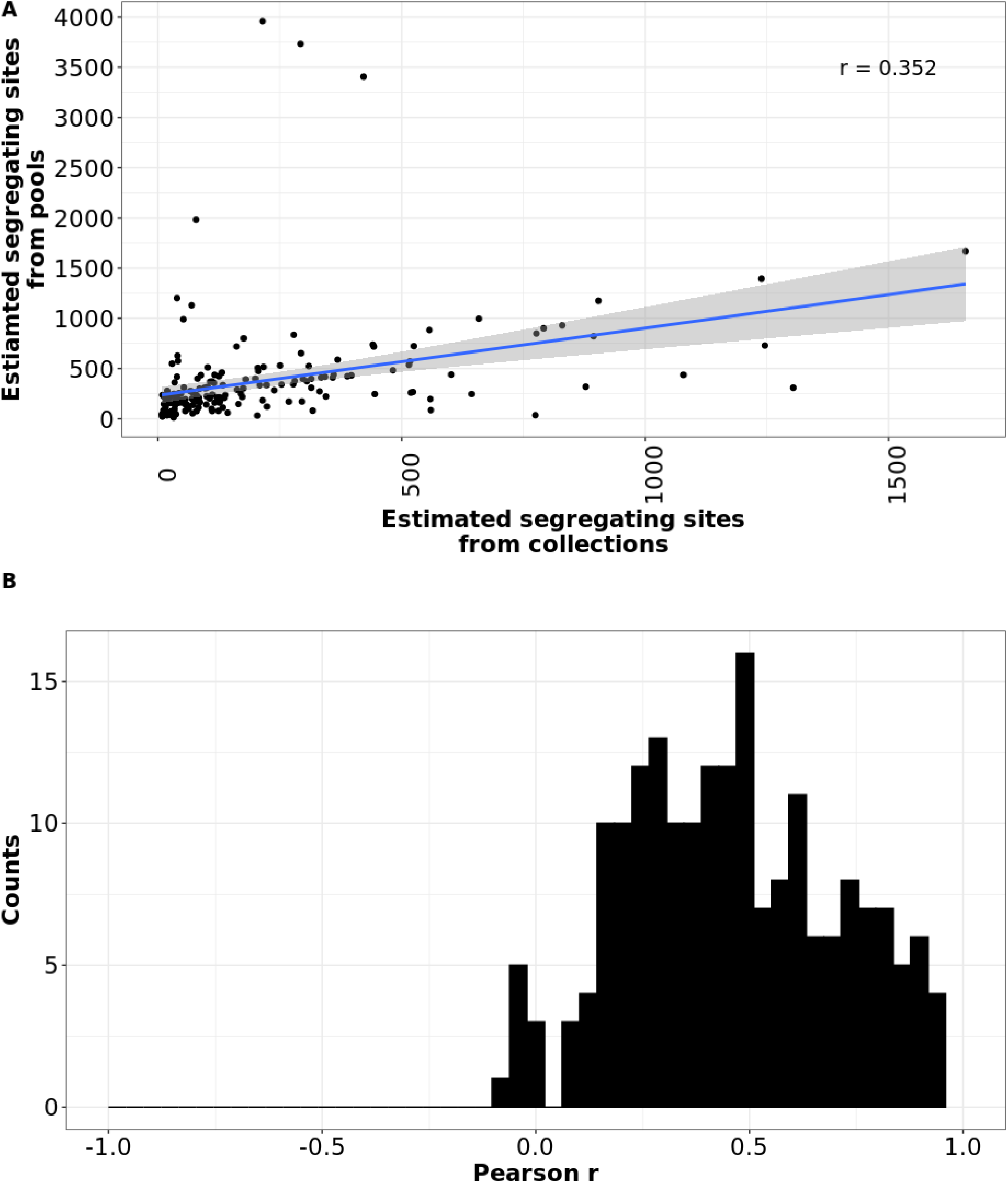
Allelic variation in pools and singles from the same sample were positively correlated. **(A) The number of segregating sites in the true pools were proportional to the number of segregating sites in the expected pool.** For mono-ST collections (collections where all eight singles, the pool and the auto-chosen reference were called the same ST), the number of sites with allelic variation was comparable between the true pools (y-axis) and the expected pool (x-axis) (eight singles combined). If the same site was fixed in all eight singles and in the pool, it was not included. Blue regression line depicts a linear relationship with a Pearson’s *r* of 0.352. **(B) AFs of variants in the expected pool did not reliably predict the AFs of the same variants in the true pool**. Frequency distribution plot showing Pearson’s *r* for all 198 mono-ST collections. x-axis depicts Pearson’s *r* and y*-* axis depicts number of collections.

### A median of one more AMR gene was detected in the pools compared to singles

Finally, we wanted to know if the pools could harbour subpopulations with clinically relevant genes that may be missing in singles. We annotated AMR genes using AMRFinderPlus and counted the number of antimicrobial drug classes for which resistance determinants were found in our pools, individual singles, and in the pangenome of our expected (all genes eight singles combined) and downsampled (all genes from two or four random singles combined) pools (36). In 177 collections (79%), the number of AMR classes was identical in pools and the expected pool. This group represented the bulk of the low-diversity samples in the study. However, overall, we observed a median of one additional AMR class in our true pools compared to the expected/downsampled pools and singles (**Fig 7, black vertical line**). This showed that additional genes could be detected in the pool that are absent in the pangenome of the singles and that these genes can be of clinical relevance. Notably, all cases where we found *mecA* in the pools (134 out of 226 pools), we found *mecA* in at least one of the eight corresponding singles. A summary file with all detected resistant determinants for all collections is reported in the supplemental file ‘**Supplemental_dataset_1.xlsx**’.

We initially used the total number of genes and the number of AMR classes in the pools as predictor variables in our logistic regression model in **Fig S1**. We found that the number of genes was highly multicollinear with CheckM contamination (VIF > 5) and the number of AMR classes had no predictive power (near identical AUC, Accuracy, sensitivity, and specificity with or without the AMR class parameter), and therefore did not include them in our final analysis.

Next, we wanted to measure the abundance of AMR genes present in both pools and the pangenome of the singles compared to AMR genes present in the pools alone. We hypothesised that in cases where an AMR gene was found in the pools but absent in the singles, the AMR gene was present at low abundances. To test this, we used Salmon (37) to estimate the abundance of AMR genes found in both pools and singles, and compared the abundances to when it was found in the pools alone [**Fig S3**]. We found that the mean copy number of genes belonging to a particular class of antibiotic was significantly lower when found only in the pools for eight out of nine AMR classes (Welch’s t test with Bonferroni correction).

**Fig 7:**
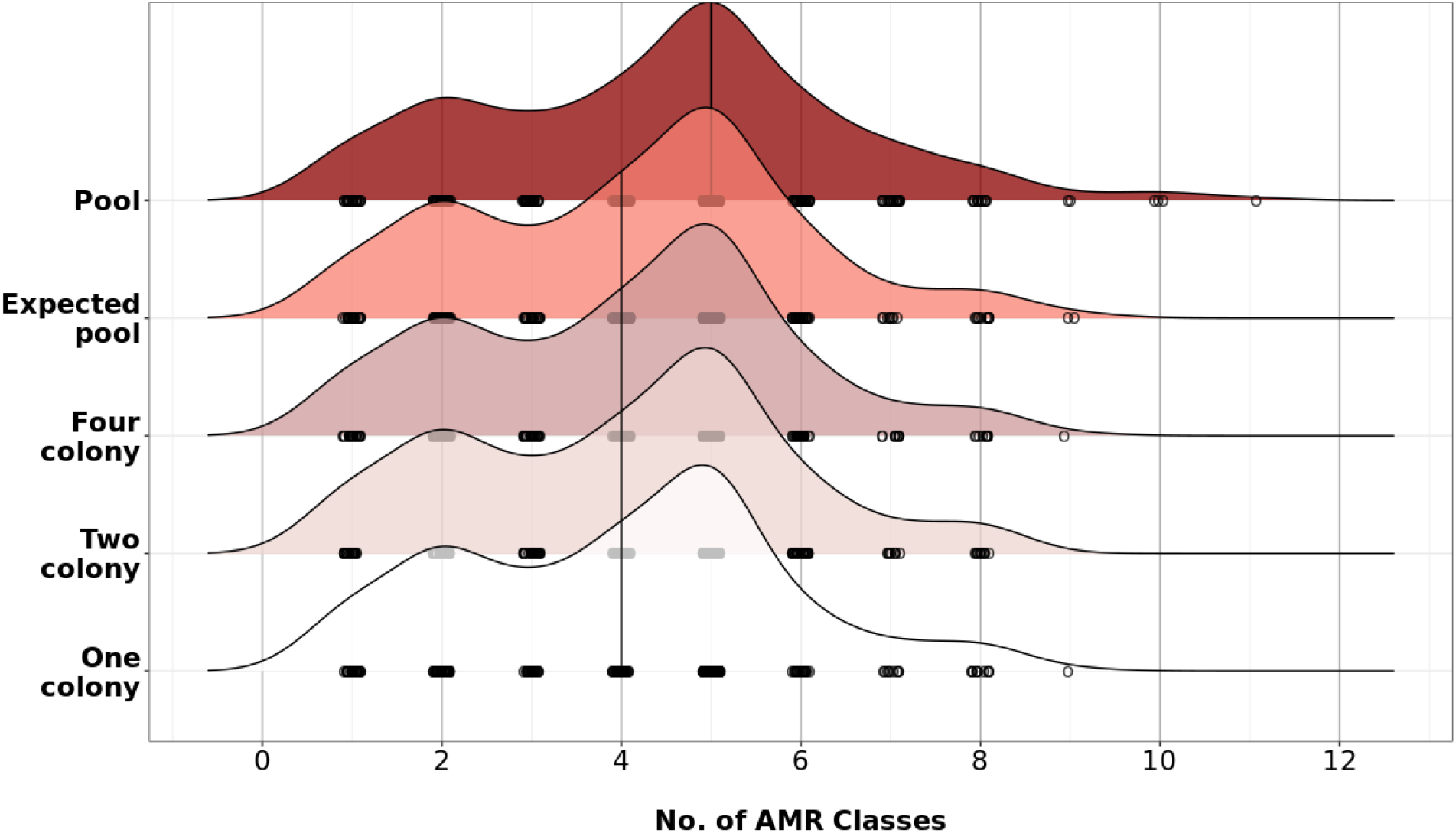
A median of one additional AMR class can be observed in the pools compared to singles. Ridgeline plot showing number of AMR gene classes detected in pools, the pangenome of expected and downsampled pools (pangenome of eight, four and two singles combined), and a random single colony. The x-axis shows the number of AMR classes detected in the sample by AMRFinder and the y-axis shows the corresponding sample. Black vertical line shows the median number of AMR classes detected for each sample group. White circles under each ridgeline represent individual collections and the number of AMR classes detected.

**Fig S3:**
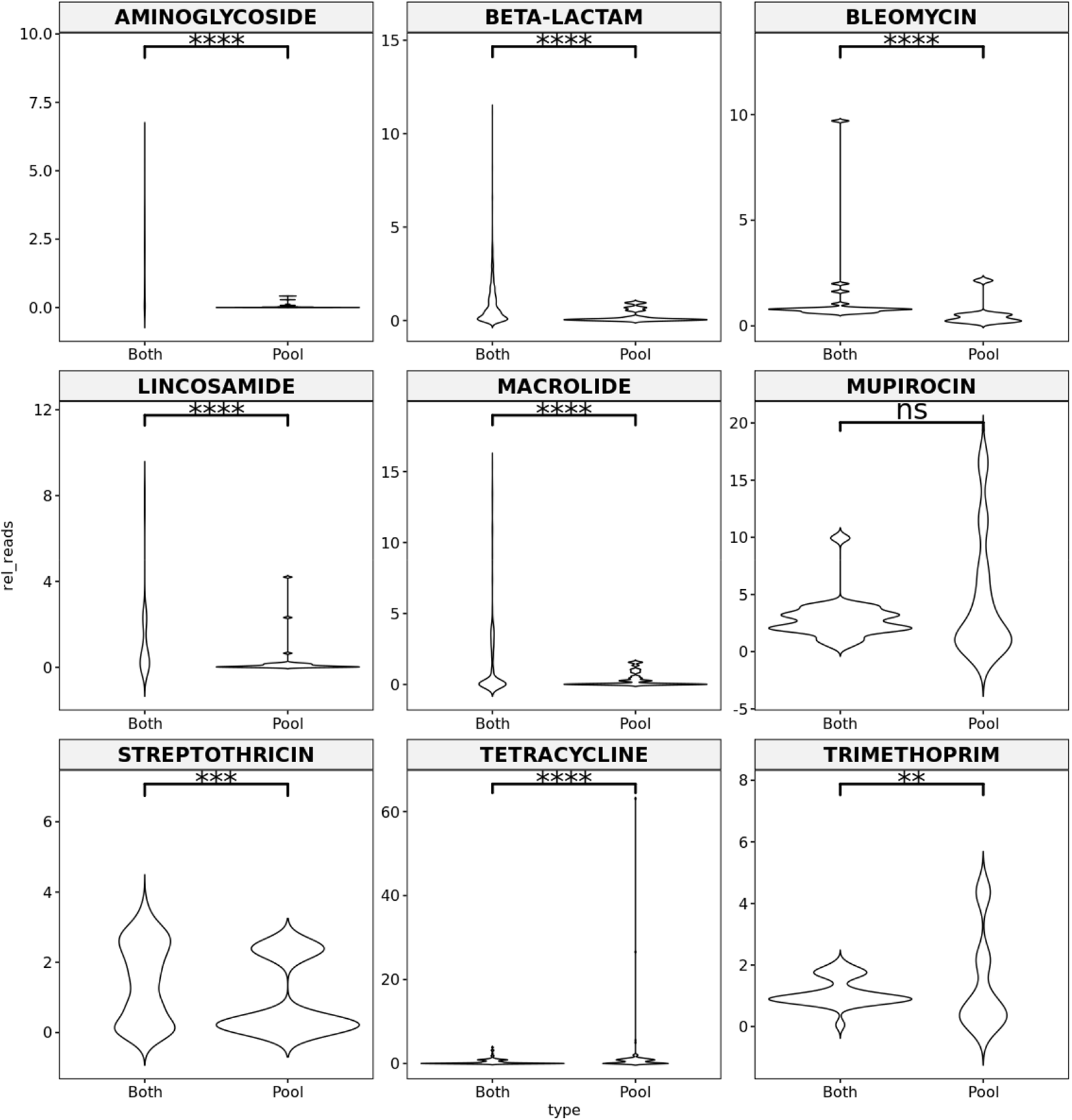
Mean read abundance is lower for AMR genes present only in the pools compared to AMR genes present in both pools and singles. For each class of AMR, we estimated the number of reads mapped to each AMR gene in the AMRFinder database relative to the number of reads mapped to *rpoD* (relative copy number). All genes were normalised to 1 kb. For each AMR class, the relative copy number of genes found in both the pool and the corresponding single (“Both”) were compared against genes for the same AMR class found only in the pool (“Pool”) using Wilcoxon rank sum test with Bonferroni correction. ns = p > 0.01; ** = p < 0.001; *** = p < 0.0001; **** = p < 0.00001.

## Discussion

From this study, we derived insights into strategies for sampling genomic diversity of *S. aureus* from asymptomatically colonised human skin and mucosal surfaces. We found that in most cases (83%), *S. aureus* populations were clonal, representing only one ST. Interestingly, there were no significant differences in the incidence of multi-ST populations across the three anatomic sites sampled (anterior nares, oropharynx (throat), inguinal skin), four different timepoints (at participant enrollment, or at three months, six months, nine months and twelve months after enrollment), nor across different culturing methods (direct vs enrichment, see methods). Many of the conclusions learned from this study could be applied by sampling other bacterial pathogens (or *S. aureus* on other hosts/anatomic sites).

The primary purpose of this study was to compare sampling strategies for measuring intra-species diversity at different levels of resolution. We wanted to understand the cost-benefit trade-offs of sampling one or multiple single colonies per sample, and pooled populations. To do this, we compared pure single colonies, *in-silico* single-colony mixtures (expected pools and downsampled pools), as well as total pools of 10s – 100s of colonies (**Fig 1**). One significant finding was that in our samples, the *S. aureus* within-host diversity for a given body site and time point was relatively low – most (83%) collections were of a single clonal lineage (**Fig 2**). Adding additional single colony genomes may be useful for certain studies focused on within-host variation and increases the chances of detecting strain mixtures but there are diminishing returns for sampling new genetic diversity within homogeneous populations (**Fig 5A**).

**Fig 8:**
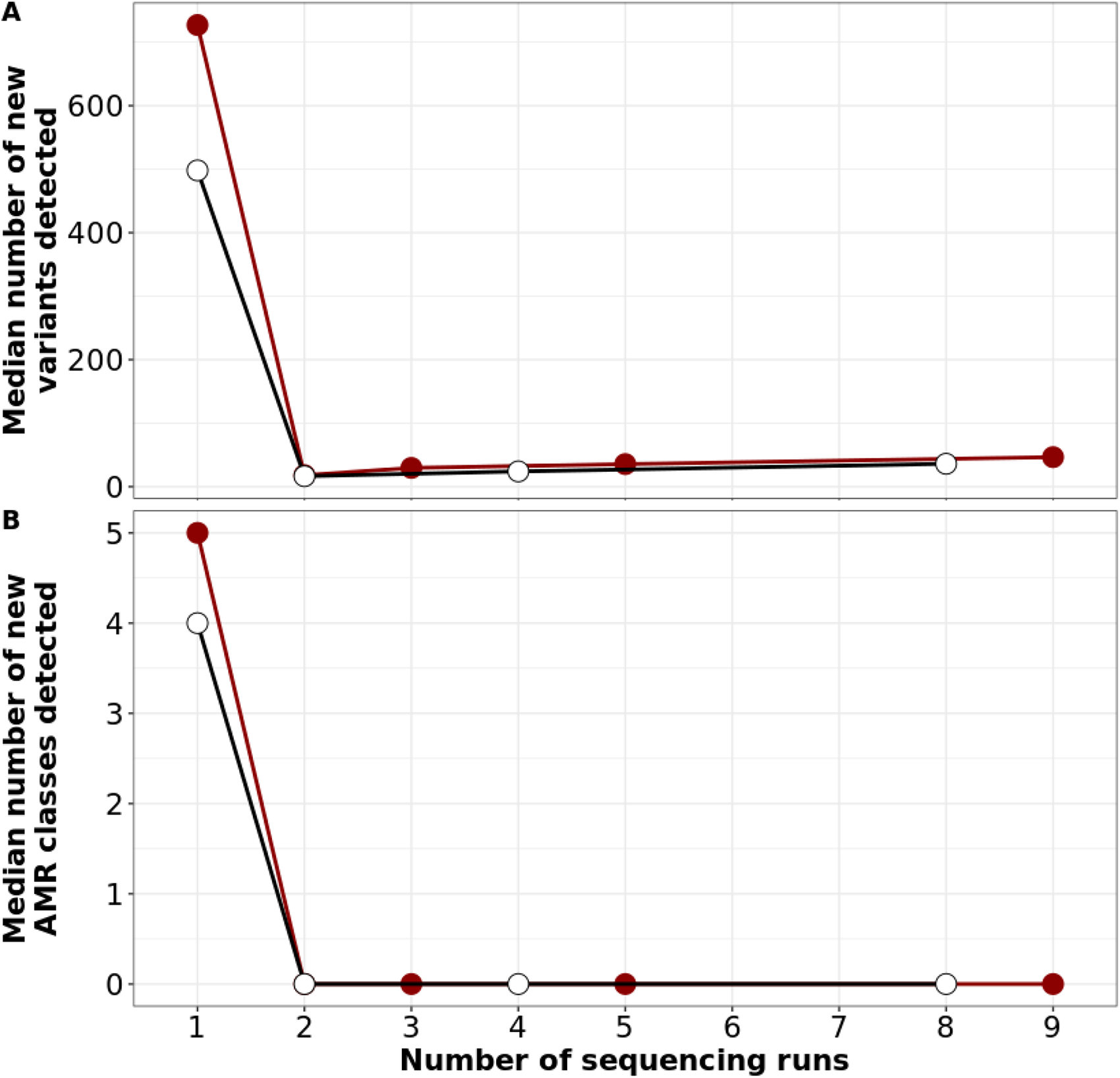
Diminishing returns in the number of new variants or new AMR genes observed with the addition of more sequencing runs. Dot plot depicting the number of new variants (**A**) or new AMR genes (**B**) observed for additional sequencing runs. Red dots depict the first sequencing run being the pool, and the additional runs being single colonies (1 = Pool, 2 = Pool + one single, 3 = Pool + two singles…). White dots depict only singles (1 = one single, 2 = two singles, …)

Assuming sequencing one pool incurs one unit cost (cost of time, labour and resources for sample preparation, storage, sequencing, and analysis), every single colony added on top of the pool would incur an additional unit cost. For our dataset, this additional unit cost over the pool yielded a median of 19 new variants and 0 new AMR gene classes (**Fig 8**). Moreover, we also showed that the pool alone is sufficient to predict the presence of a multi-ST population (**Fig 3,4, S1**).

The study had some limitations. First, the sampling methodology may in some cases have resulted in reduced diversity in the total pool by picking singles prior to pooling the remaining colonies. Our assumption in the design was that the population in a single colony may be present multiple times on a plate with 100s of colonies, but this is less likely to hold true in scenarios where there were < 15 – 20 colonies left after picking singles. There are other limitations that are inherent with incorporating culture steps. Ideally, the number of colonies in a pool should be relatively constant across all samples but in some cases very few colonies appeared on the selective agar. In other cases, the density of colonies was too great to measure colony counts accurately. Moreover, laboratory culture media could also cause biases in population growth.

Second, our sampling space is narrow – all our samples are from one geographical location, comprising only nares, throat, and skin swab-acquired *S. aureus*. Therefore, the diversity we measured across singles and pools from a given sample may not apply to cultures from other clinical contexts of *S. aureus* or other *S. aureus* strain types typically colonising people worldwide. Despite these drawbacks, the data in this study allowed us to compare three strategies for sampling: individual single colonies, collections of up to eight colonies, and pool-seq.

Sampling single colonies in pure culture is the traditional approach for assessing the genotypic and phenotypic characteristics of bacterial pathogens. Only one sequencing library is required and the bioinformatic analysis methods are straightforward. However, sampling only one colony will result in missing multi-strain infections (17% of the time in our case) and therefore provide an overly simplistic view of the population structure of the pathogen.

The more single colonies sequenced, the better the estimation of true population diversity (assuming there are no systematic sampling biases in how colonies are picked from the culture plate). Having collections of single colonies also allows the construction of within-host phylogenies, observing gene gain/loss events and inferring demographic changes over longitudinal sampling. However, there is still no guarantee that the total diversity in the population is represented in the sample subset. Moreover, the cost of processing and physical and data storage scales linearly with the number of independent colonies sampled. Deciding the number of colonies to sample will be a complex calculus of budget and *a priori* estimation of the population diversity of the pathogen aligned with the goals of the study.

Ideally, pool-seq would provide the best estimation diversity in the population, and many more single colonies can be aggregated than sequenced individually. After pooling colonies, the stock can be treated as a single sample for storage, sequencing, and analysis; therefore, the cost is equal to one single colony, making pool-seq the best value for identifying variation. Here, we have shown that pool-seq can be used to accurately estimate the presence of multi-ST infections, can measure segregating sites within mono-ST populations, and are most sensitive at finding AMR genes. The analysis of pool-seq data, which are effectively single-species metagenomes, is more complex than single colony sequencing due to variation, especially in multi-ST samples, and the possibility of contamination. This heterogeneity also leads to unreliable phenotypic ascertainment. However, predicted phenotypes could be validated by replating the pool to pick singles.

Based on our analysis, we recommend that many studies may benefit from using a **one plus pool** design, that is, sequencing one single colony + pooling and sequencing all remaining colonies. Using the sample diversity measurements we show in this study, many of which are obtained from the default outputs of Bactopia, a streamlined beginner-friendly analysis pipeline, we can ascertain whether significant differences exist between the single colony and the pool (**Fig 3,4**). This can aid in deciding whether sequencing additional colonies from the pool is required. We believe this approach provides more information than a single colony while demanding little additional time and labour for sample collection, storage, and analysis. A disadvantage of processing pooled samples is the reduced sequence quality and increased likelihood of contamination (**Fig 3**). However this disadvantage can be mitigated with the additional sequencing of at least one pure single colony.

## Methods

### Strain sampling

Participants were enrolled into the SEMAPHORE study after presenting with a *S. aureus* positive SSTI. Up to four timepoints (every three months for one year) and up to three body sites (Anterior nares, oropharynx, and inguinal skin) were sampled for each participant. Each swab was streaked out onto BBL^TM^ CHROMAgar^TM^ *Staphylococcus aureus* (SACA). Then eight individual colonies (singles) were subcultured onto blood agar and sequenced. The remaining colonies were pooled, sub-cultured onto blood agar, then sequenced. In cases where there was no growth from directly plating the swabs, each swab was enriched for growth in tryptic soy broth (TSB) overnight. These overnight cultures were then plated, and colonies were picked as mentioned above. 181 pools and 1448 singles were directly plated on SACA (Direct cultures).

The remaining 73 pools and 584 singles did not show growth upon direct plating and therefore were enriched in TSB and then plated (enrichment cultures). In this study, we only analysed swabs from which eight singles and a pool were obtained. Swabs from which fewer than eight singles were obtained were not considered. In total, we obtained 254 pools from 85 participants and eight singles corresponding to each pool, giving us a total of 2286 genome sequences.

### Library preparation and sequencing

Genomic DNA extractions were performed using Qiagen kits. Library preparation and whole genome sequencing were performed by the Children’s Hospital of Pennsylvania Microbiome Center using Illumina MiSeq or Hiseq platforms.

### Genome assembly, annotation and variant calling using Bactopia

All obtained sequences were processed using the Bactopia analysis pipeline v1.75 (29). Bactopia performed adapter trimming using BBTools (38), genome assembly using SKESA (30) and the assembly quality was assessed using QUAST and CheckM (31,32). Genome annotation was done using Prokka (39) and AMR genes were annotated using AMRFinderPlus (36). Variant calling was performed by Snippy (40) using an automatically selected reference sequence based on the closest MASH (41) distance to a complete *S. aureus* genome sequence in NCBI RefSeq. MLST types were identified using MLST (42).

### Pairwise SNP distance calculation, dereplication, and phylogeny

For each group of eight singles in a collection, we used Parsnp v1.5.3 to align the single colony isolated genomes and used snp-dists v0.7.0 to calculate pairwise SNP distances (43). To dereplicate singles, isolates with SNP distance < 10 were collapsed into clusters and a random isolate was chosen as the cluster representative using Assembly-Dereplicator (44). The final set comprised 294 singles, where each collection is represented at least once. This resulting set of singles was aligned again using parsnp and a core genome phylogeny was constructed using FastTree (45,46). Phylogeny was visualised using ggtree (47)

### Number of variants, segregating sites, and allele frequency calculation

We calculated allele frequencies from bam files generated by Bactopia using bcftools mpileup (48). The reference for each pool was auto-chosen by Bactopia based on the closest complete *S. aureus* genome in terms of MASH distance (41). All singles were then aligned to the same reference as their corresponding pool. Only variants with a QUAL score > 50 and with at least a read depth of 25 were considered for the analysis. For each collection, we calculated the allele frequencies for every position across the genome where at least one read piled up with a base call differing from the reference allele. Variants with frequencies < 0.05 were considered 0 (absence) and > 0.95 were considered 1 (fixed). The allele frequencies for the expected and downsampled pools were calculated based on the number of singles the variant was fixed in. For example, if a variant was present in one out of the eight singles at a frequency > 0.95 (fixed), its allele frequency in the expected pool would be or 0.125. Two or four random singles out of the eight ⅛ were selected to measure the allele frequencies in the downsampled pools. Variants with intermediate frequencies in the singles (>0.05 & < 0.95) were not considered.

To calculate the number of segregating sites across the true and expected pools, we wanted to exclude variants that occurred simply because of alignment against a specific reference. If a given variant was fixed in the expected pool (present in eight out of eight singles at an AF > 0.95) and in the true pool (AF > 0.95), we considered these variants to be ancestral and did not count them as segregating sites. All remaining sites with AF > 0.05 were counted.

We calculated nucleotide diversity (π) using InStrain (33) using the auto-chosen reference and π the alignment bam file from Bactopia. Expected pools (eight colonies) and downsampled pools (two and four colonies) were generated by combining equal proportions of reads from all eight, two or four colonies. For each collection, we used reformat.sh from the bbtools suite (38) to sample reads 12.5% from all eight colonies for the expected pool, 50% of reads from two randomly selected colonies for the two-colony downsampled pool, and 25% of reads from four randomly selected colonies for the four-colony downsampled pool. All artificial pools (expected and downsampled) contained 1 million reads.

### Logistic regression

Logistic regression was performed in R using the glm function (49). 70% of 254 pool-seq samples were used as the training set and the remaining 30% was used as the test set. A pool was considered multi-ST if the MLST alleles in the pool and the corresponding eight singles were not identical. Continuous probabilities from the logistic regression model were converted to binary using a cutoff of 0.89 (If probability > 0.89, the prediction was considered to be multi-ST). This cutoff was estimated using the optimalCutoff function from the R package InformationValue (50). McFadden R^2^ was calculated using the pR2 function from the R package pscl (51). Variance Inflation Factor was calculated using the vif function from the R package car (52).

### Statistical analyses and data visualisation

All statistics and PCA were performed in R using packages stats and rstatix (49,53). All plots were visualised using R package ggplot2 (54). Other graphics were created using bioicons and draw.io (55,56).

## Data availability

All code and raw data are available at https://github.com/VishnuRaghuram94/GASP. All genome sequences used in this study are available under PRJNA918392.

## Acknowledgements

TDR and MZD acknowledge funding from NIH NIAID 1 R01 AI158452-01A1. VR was supported by NIH AI139188 and by Emory University and the Infectious Disease Across Scales Training Program (IDASTP). We thank Dr. Joanna Goldberg for providing constructive comments on the manuscript.

## Data summary

Genomes sequenced for this study are available under accession PRJNA918392. Raw data and code for analysis are available at https://github.com/VishnuRaghuram94/GASP.

## Impact statement

While pooled population sequencing has been employed to study within-host diversity, the differences in attainable information between single and pooled sequences are not clear. A direct comparison between single isolate and pooled population sequences can help devise optimal sampling strategies for clinical and within-host diversity studies. In this study, we attempt to answer the question of how many colonies obtained from a single patient is enough to obtain a representation of the total population diversity within the patient while keeping in mind time and labour costs. These findings have implications for using whole genome sequencing (WGS) in the clinical microbiology laboratory to identify and speciate pathogens and to determine their antimicrobial susceptibilities.

